# Blood sodium regulation by Na,K-ATPase: Euryhaline animals’ salinity adaptations described by the pump’s negative feedback

**DOI:** 10.64898/2026.01.08.698424

**Authors:** Peter Ruoff

## Abstract

The term enantiostasis was introduced by Mangum and Towle to describe a functional relationship between environmental changes and an organism’s adaptation, and furthermore, to distinguish this kind of adaptation from homeostatic control. The defining example of enantiostasis is the blue crab’s blood salinity adaptation when the animal is exposed to different environmental salinities during its move between the ocean and upriver locations. In this paper a model for osmoregulation is suggested based on one of two identified negative feedbacks in the mechanism of Na,K-ATPase. Using a scheme suggested by Towle in 1997 the effect of the feedback is tested with respect to the blue crab’s salinity adaptation. Not only does this single feedback model give a practically quantitative description of the blue crab’s adaptation data, the model is also able to show other observed patterns of osmoregulation. In view of these results the relationship between enantiostasis and homeostasis is discussed.

**Graphical Abstract:** 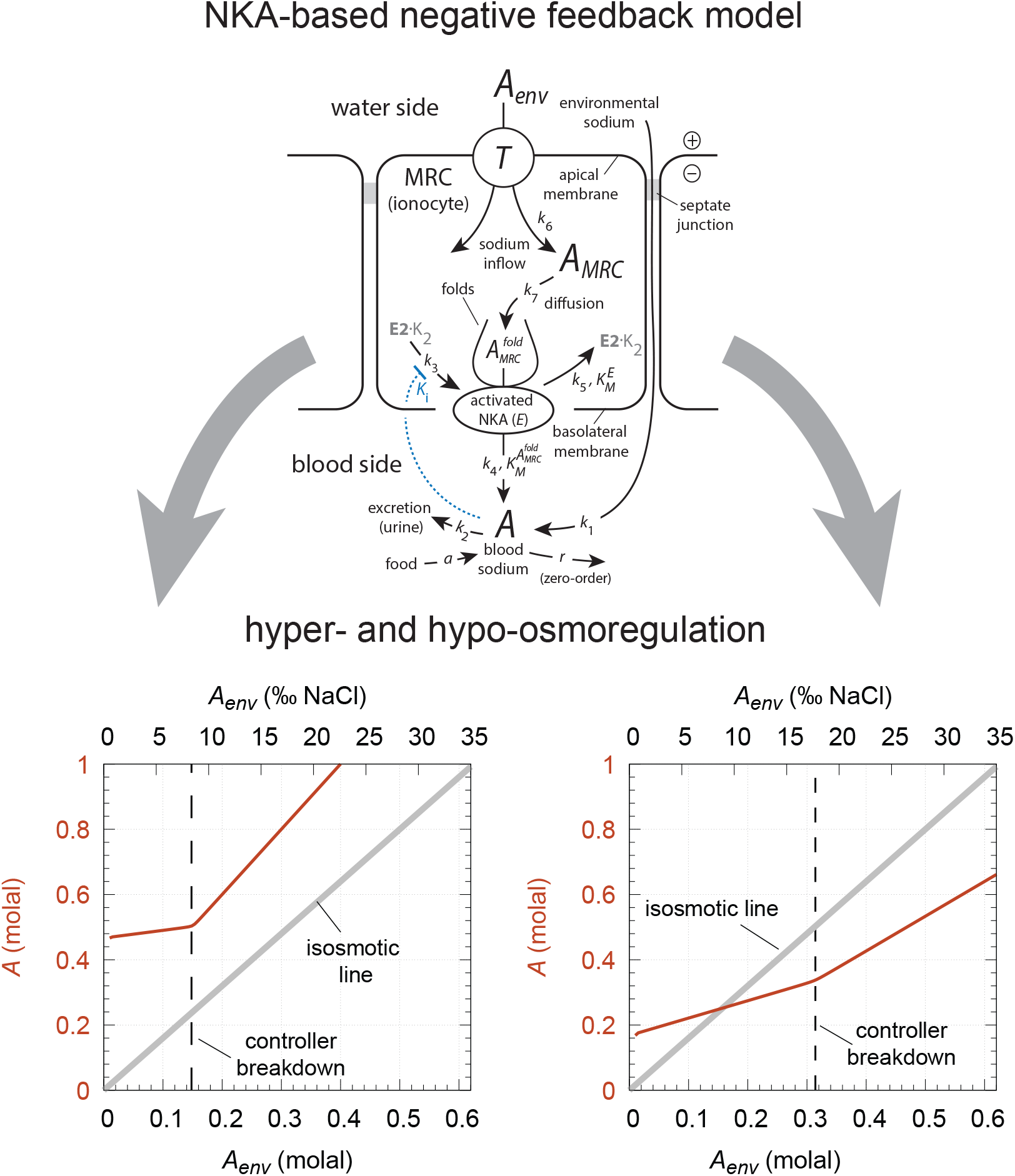

**Highlights:**

- Two sodium-regulating feedbacks are identified in the operation of Na,K-ATPase.
- One of these feedbacks is able to describe hypo- and hyperosmotic regulations.
- The relationship between enantiostasis and homeostasis is discussed.

## 1. Introduction: The concepts of homeostasis and enantiostasis

One of organisms’ essential features is their ability to adapt to changed environmental and anticipated conditions. Homeostasis, a concept coined by Walter Cannon in 1929, is a central adaptation strategy by keeping most of an organism’s steady states within relative narrow limits (Cannon, 1929; Langley, 1973; Clancy and McVicar, 2002). However, other replacing concepts were later suggested. Mangum and Towle (1977) introduced the term *entantiostasis* to highlight an adaptation strategy with a *conserved functional relationship* between a changing environment and the organism’s internal steady states (Hochachka and Somero, 1984).

The defining example of enantiostasis is shown in Fig 1a, which illustrates how blood osmolality in the blue crab *Callinectes sapidus* changes as the animal encounters different environmental salinities ranging from saltwater (35 ‰) to fresh water (Mangum and Towle, 1977). Since the slope in blood osmolality between 1 ‰ to about 27 ‰ is significantly greater than zero, Mangum and Towle concluded that this type of adaptation is opposite to homeostatic and termed it enantiostatic. In fact, the increased slope could also be characterized as a *rheostatic* response, a concept which was introduced by Mrosovsky (1990) focussing on setpoint changes (Stevenson, 2024), but apparently without being aware of the close similarities between these two approaches. An interesting feature in osmoregulation is the appearance of a “breakpoint” (Fig 1a), which often occurs at higher environmental salinities.

**Figure 1:**
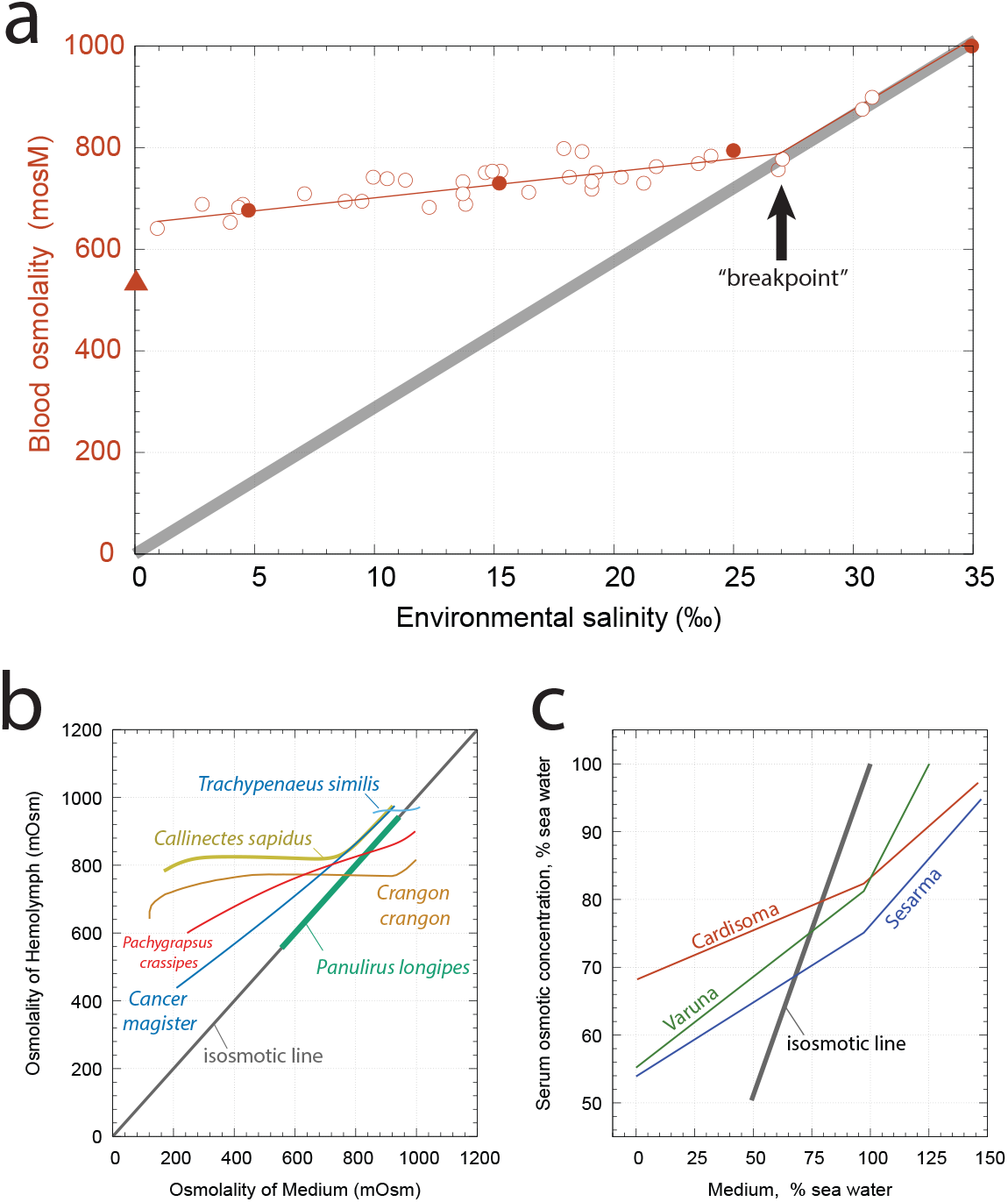
Types of osmoregulation. Panel a: Blood osmolality as a function of environmental salinity in the blue crab *Callinectes sapidus*. Open circles represent values of caught animals in various salinities. The triangle is the average of crabs caught in freshwater. Solid dots represent mean values of crabs acclimated at different salinities in the laboratory. The grey thick diagonal is the isosmotic line. It describes the situation when internal and environmental osmolalities are in equilibrium. Redrawn from Fig 2 of Mangum and Towle (1977). Panel b: Scheme of other hyperregulators including the osmoconformer *Panulirus longipes*. Redrawn from Figure 1 of Mantel and Farmer (1983). Panel c: Examples of hyposmotic regulators. Redrawn from Figure 1-15 of Prosser (1973).

Mangum and Towle (1977) ascribed the blue crab’s hyperosmotic behavior in Fig 1a (i.e. blood osmolality is higher than the osmolality of the surroundings) to a synergistic interplay between osmoregulation by Na,K-ATPase (NKA), oxygen transport, and nitrogen metabolism. The authors argued that a homeostatic, i.e. unchanged, blood salt content does not seem possible, because when crabs are moved from higher to lower water salinities the decrease in the blood’s osmolality is a trigger for the production of counterions to drive the salt pump (Mangum and Towle, 1977). In addition, many studies have shown that maintaining hyperosmoregulation in crabs is accompanied by an upregulation of the sodium pump when crabs are transferred from a high salinity environment to a lower one (Towle et al., 2001). On the other hand, when testing NKA’s *α*-subunit mRNA and protein in crabs moved from a 35 ‰ salinity to 5 ‰ no notable differences in mRNA or protein were observed. This led Towle et al. (2001) to the conclusion that the hyperosmoregulatory response in the blue crab may result from post-translational regulatory processes.

Fig 1b shows responses of other hyperosmoregulators, which differ from the blue crab both in slopes as well as where the “breakpoint” occurs. The osmoconformer *Panulirus longipes* is included, which keeps its blood osmolality at the isosmotic line where osmolalities of blood and seawater are equal. In Fig 1c examples of hyposmotic regulators are shown. These animals are normally more dilute in their body fluids than sea water and live in hypersaline lakes (Prosser, 1973).

### 1.1. Mechanism of NKA and dual feedback structure

NKA follows a ping-pong scheme termed as the Albers-Post mechanism (Albers, 1967; Post et al., 1969; Jorgensen et al., 2003; Dyla et al., 2020) (Fig 2a). In this scheme NKA cycles between two conformational states, *E*1 and *E*2, where potassium bound NKA in the *E*2 state binds ATP and phosphorylates the enzyme. This state releases 2 potassium ions into the internal space, thereby switching to the *E*1 conformation which binds 3 sodium ions. As pointed out by Skou and Esmann (1992) and Skou (1998), a characteristic feature of NKA is that the phosphorylation by ATP is activated by a combined effect of Na^+^ on the cytoplasmic side and K^+^ on the extracellular side and induces a conformation change. Skou and Esmann (1992) stress further that experiments performed on purified NKA reconstituted into liposomes show that the ATPase activity not just represents the catalytic part, but also contains the translocating part of these cations (Goldin, 1977; Cornelius and Skou, 1988). Panel b shows the dual feedback structure of the sodium pump. One of the feedbacks is based on a concentration dependent activation of NKA by internal sodium ions (Cornelius and Skou, 1988; Skou and Esmann, 1992) with the subsequent sodium expulsion by the pump (outlined in red). The other feedback occurs due to the inhibition of the pump by external sodium (Garrahan and Glynn, 1967; Sachs, 1977; Kaplan, 1982) (outlined in blue). It is the inhibitory feedback, which we will investigate.

**Figure 2:**
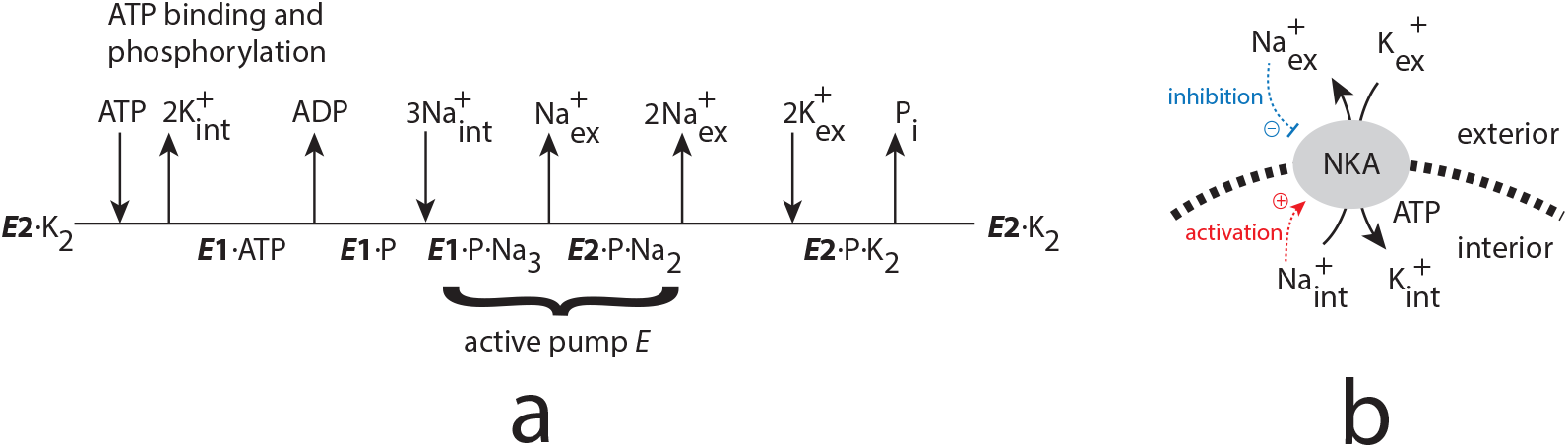
Mechanism of the sodium pump and the pump’s dual feedback structure. Panel a: Cleland notation (Cleland, 1963) of the Albers-Post mechanism (after Jorgensen et al. (2003)). The two conformations, which pump sodium ions are indicated. Panel b: The activation of NKA by internal sodium (outlined in red) and its inhibition by external sodium (outlined in blue) create in both cases a negative feedback with respect to internal or external sodium.

## 2. Materials and methods

### 2.1. Calculations and graphical presentations

Computations were performed with the Fortran subroutine LSODE (Radhakrishnan and Hindmarsh, 1993), which can be downloaded from https://computing.llnl.gov/projects/odepack. Gnuplot (www.gnuplot.info) was used to generate figures, while reaction schemes and plot annotations were prepared with Adobe Illustrator (www.adobe.com). A set of Python scripts using LSODA are provided for documentation (see Supporting Material).

### 2.2. The model

As remarked by Mantel and Farmer (1983) the presence of up to 10% protein in the blood (hemolymph) of euryhaline animals makes a direct comparison of ionic compositions between media difficult. In the model the presence of protein and other ions is ignored by taking blood osmolality as sodium osmolality. Since sodium chloride is a dominating component in seawater the environmental salinity in Fig 1 is replaced in the model by sodium molality and ‰ NaCl. This makes model data only approximately comparable with experimental data, but still addresses the essential feature of osmoregulation by the sodium pump.

The model, shown in Fig 3, is based on the transport system described by Towle (1997) (see Fig 3 there). Instead of the three different transporters by which environmental sodium (*A*_*env*_) enters the mitochondrial rich epithelial cells (MRCs, ionocytes), a generic transporter *T* is used. For simplicity, sodium enters the cytosol of the MRCs as a first-order reaction with respect to *A*_*env*_ with rate constant *k*_6_ and velocity *v*=*k*_6_·*A*_*env*_.

**Figure 3:**
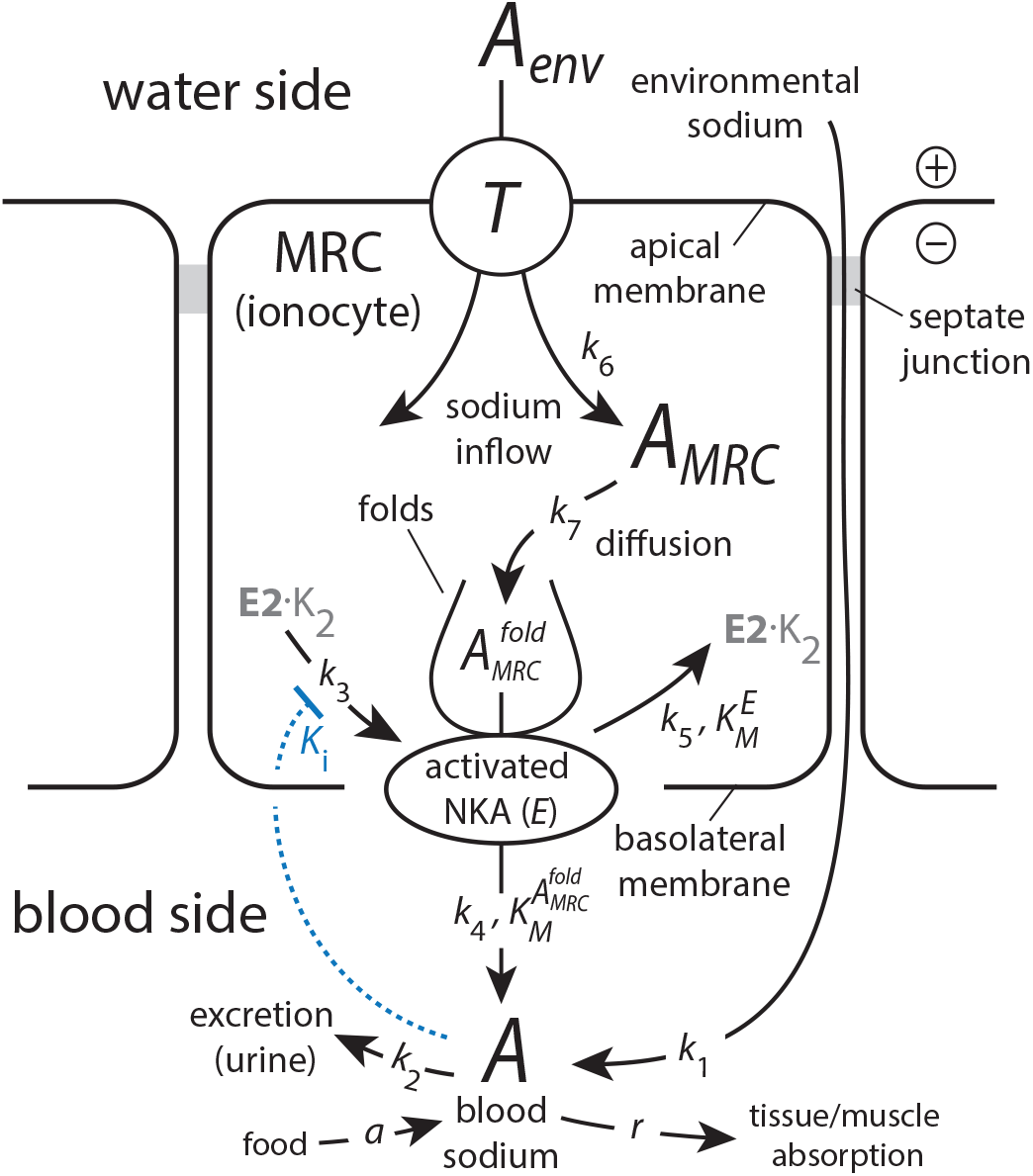
Negative feedback model of blood sodium homeostasis in euryhaline animals. Environmental sodium, *A*_*env*_, enters mitochondrion rich cells (MRCs) via several transporters (Towle, 1997), lumped here together as *T*. Another entry route directly into the blood is considered to occur through septate junctions. Sodium ion inside MRC, *A*_*MRC*_, diffuses to the basolateral side of the cell where NKA is situated inside a folding structure. Sodium ions located inside these folds, 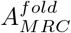, are transported by activated NKA (variable *E*) into the blood. Activated NKA is produced from the *E*2 · *K*_2_ state (Fig 2a) and recycled to it. The negative feedback occurs by inhibiting the activation of NKA by blood sodium (variable *A*) with inhibition constant *K*_*i*_ (outlined in blue). For rate equations, see main text.

Sodium inside the MRCs, *A*_*MRC*_, diffuses to the basolateral membrane where NKA is located (Towle and Kays, 1986). The membrane structure there shows invaginations or foldings which increase the interfacial area between blood and the basolateral membrane, and thereby increase the local NKA densities. As indicated in Fig 2b, *A*_*MRC*_ takes part in the activation of NKA leading to the *E*1· *P* ·*Na*_3_ and *E*2 · *P* · *Na*_3_ states. In the model these two states are lumped together as *E* (activated NKA), which moves sodium ion into the blood.

The blood sodium concentration is described as *A*. Included in the model are leaky septate junctions situated between epithelial cells. Due to the apical transporters, here formulated as *T*, an outside-positive potential is considered to build up, which can drive a substantial inward paracellular sodium influx across the leaky junctions (Luquet et al., 2002; McNamara and Faria, 2012; Henry et al., 2012). In the model this is formulated as a first-order process with rate constant *k*_1_. The excretion of *A* via the urine is described as another first-order process with respect to *A* and rate constant *k*_2_. A food-related rate *a* into the blood sodium pool is included in addition to a sodium storage term *r* which moves blood sodium into muscles and other tissues (Titze, 2015).

The rate equations are:

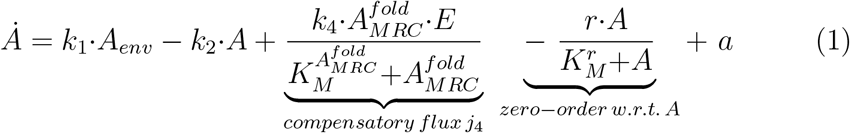

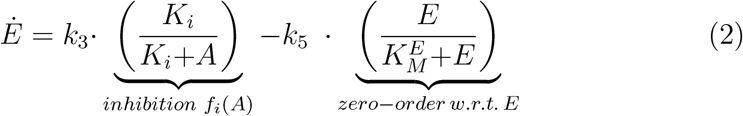

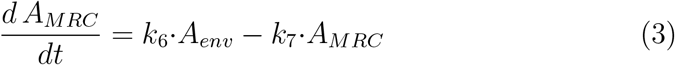

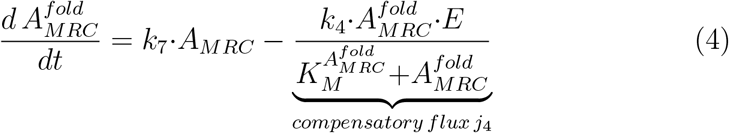

#### 2.2.1. Integral control, robust perfect adaptation, and feedback breakdown

The model has the potential to show integral control, which is a method from control engineering (Wilkie et al., 2002). Since it turns out that the model’s ability to show integral control (or close to integral control) appears to be a relevant part when describing the behavior in Fig 1, the method is briefly outlined.

Integral control or integral feedback is a method from control engineering applied with respect to a negative feedback, and which requires the presence of a setpoint of a controlled variable. The difference (error *ϵ*) between the setpoint and the actual value of the controlled variable is calculated and integrated in time. The integrated error is then fed back into the feedback. Integral feedback leads to robust perfect adaptation (Yi et al., 2000; Khammash, 2021), where the controlled variable is moved precisely to its setpoint independent of environmental perturbations.

In chemical systems integral control can be implemented by several kinetic approaches (Yi et al., 2000; Ni et al., 2009; Drengstig et al., 2012; Shoval et al., 2010; Briat et al., 2016; Fjeld et al., 2017; Aoki et al., 2019; Waheed et al., 2022). Here, the model can show integral control in *A* when the second term in Eq 2 becomes close to zero-order with respect to *E*, i.e. when 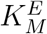 is low in comparison with *E* and 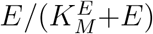 is close to one. At steady state, Eq 2 then defines *A*’s setpoint at *A*_*set*_=*K*_*i*_(*k*_3_ − *k*_5_)*/k*_5_, which can be seen by rearranging Eq 2:

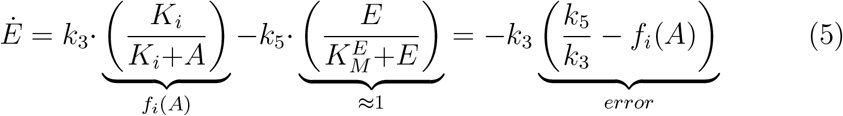

When approaching steady state the error in Eq 5 goes to zero and we get:

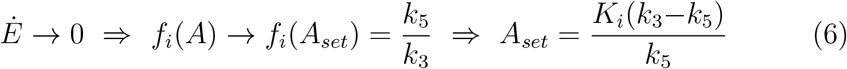

In the present feedback model the error is not a direct difference between a setpoint and the controlled variable, but between the reference value (*k*_5_*/k*_3_) and *f*_*i*_(*A*), as described by Eq 5. The setpoint of *A* is then calculated from the condition *f*_*i*_(*A*_*set*_)=*k*_5_*/k*_3_ by solving for *A*_*set*_ (Eq 6).

## 3. Results

### 3.1. Modeling the blue crab’s osmoregulation

Fig 4 shows the model’s *A* concentration as a function of *A*_*env*_. The experimental data by Mangum and Towle (Fig 1) are indicated as open squares. To increase *A* with growing *A*_*env*_ *K*_*i*_ is slightly increased with increasing *A*_*env*_, i.e. in the calculations the following relationship is used:

**Figure 4:**
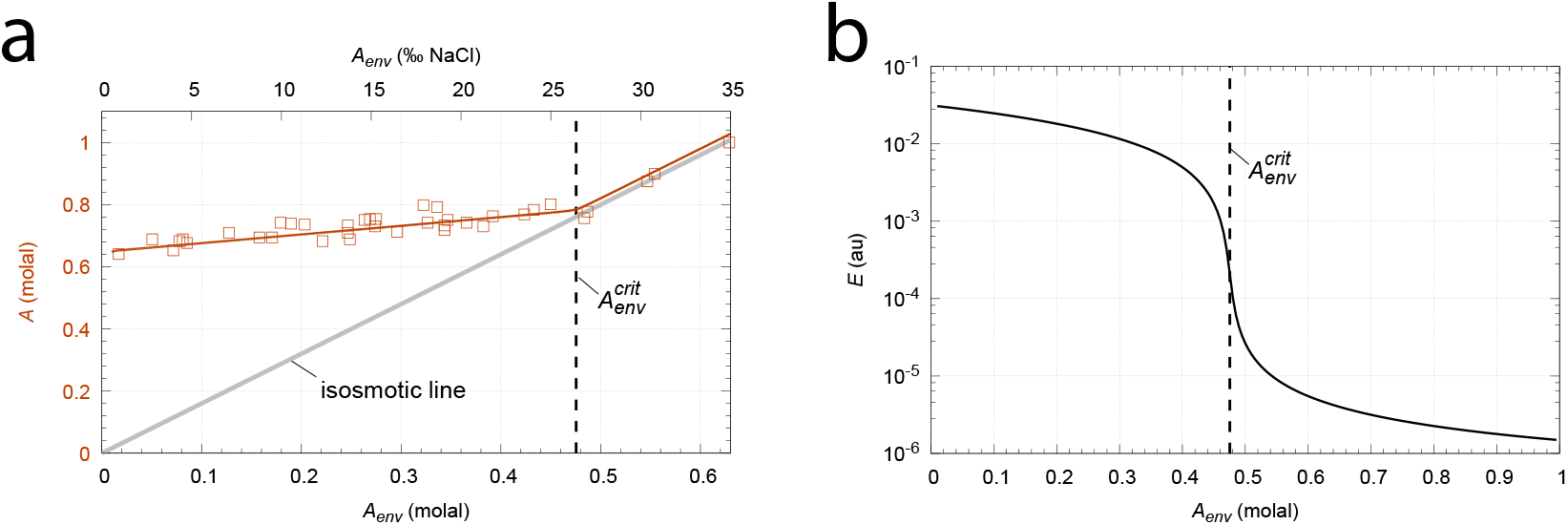
The blue crab’s enantiostatic behavior simulated by the model (Fig 3). Panel a: Concentration of blood sodium (*A*) as a function of environmental sodium concentration (*A*_*env*_). The thick grey line describes the isosmotic line. Panel b: Amount of activated NKA (*E*) as a function of *A*_*env*_. The dashed vertical lines indicate 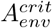 described by Eq 11. Rate constants (au): 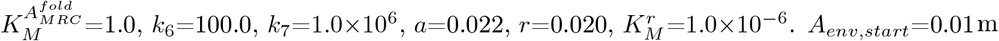 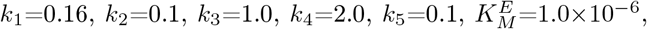 and *K*_*i,start*_=0.072 m, *A*_*env,end*_=1.0 and *K*_*i,end*_=0.103. Initial concentrations: *A*=0.6 m, *E*=0.35 au, *A*_*MRC*_=1.0 au, 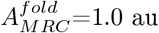. In the Fortran calculation the *A*_*env*_ range was divided into 200 data points. For each *A*_*env*_ value the steady state was determined after 2000 time units. See Fig4.py for a corresponding Python calculation.

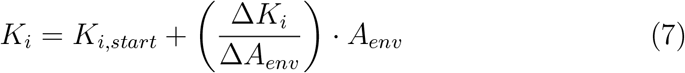

with

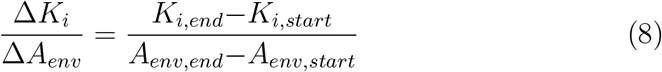

A characteristic property of the model is that, as *A*_*env*_ increases, *A* homeostasis is maintained by decreasing the compensatory flux *j*_4_ (Eq 1). Or, as crabs move towards more dilute environments blood sodium homeostasis is achieved by derepression of the pump via *A*, which causes an apparent increase of the pump’s activity. Increased *A*_*env*_ values finally lead to a critical concentration, 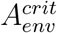, when *j*_4_ becomes zero and the controller breaks down. By using *j*_4_=0 in Eq 5 the following “breakdown line” can be defined:

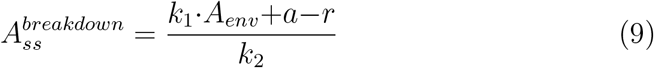

The intersection point between 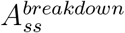 (Eq 9) and the *A*_*env*_-dependent setpoint expression Eq 10 (which is obtained by inserting Eq 7 into Eq 6)

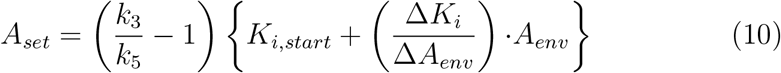

determines 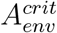 as:

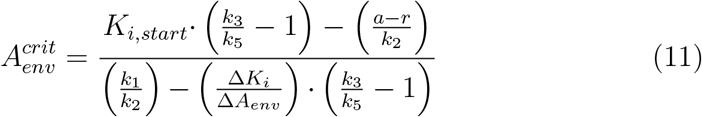

As Fig 4a indicates, it is at 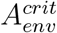 the controller breakpoint in Fig 1 occurs. As *A*_*env*_ further increases the steady state values of *A* now follows the breakdown line Eq 9, which is different from the isosmotic line.

### 3.2. Robust homeostasis, controller aggressiveness, and accuracy

When 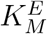 is low (zero-order conditions with respect to *E*, Eq 2) *A*_*set*_ depends only on *k*_3_, *k*_5_, and *K*_*i*_. The controller shows robust perfect adaptation, and changes in the other rate constants do not affect *A*_*set*_. This is illustrated in Fig 5 for varying *k*_1_ and *k*_2_ values during four phases when *A*_*env*_=0.2 m and *K*_*i*_=7.735 × 10^−2^ leading to a *A*_*set*_ of 0.696 m. In phase 1 *k*_1_ and *k*_2_ and the other rate constants have the same values as in Fig 4, while in phase 2 *k*_1_ is doubled. In phase 3, *k*_1_ is returned to its value during phase 1, while *k*_2_ is trippled. Finally, in phase 4 both *k*_1_ and *k*_2_ are significantly reduced. Despite these changes *A* is kept robustly at *A*_*set*_=0.696.

**Figure 5:**
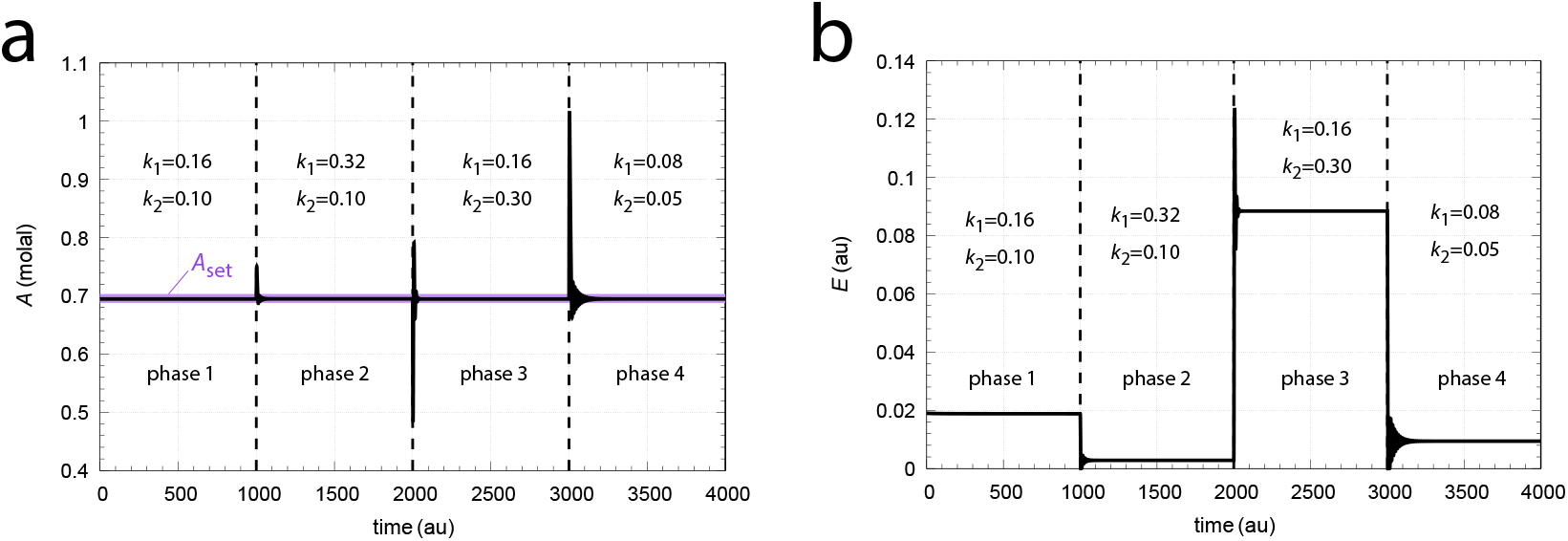
Perfect adaptation of *A* at *A*_*set*_=0.696 (violet line) when *k*_1_ and *k*_2_ are changed and *A*_*env*_ is constant. Panel a: *A* as a function of time. Panel b: *E* as a function of time showing its perturbation dependent repression and derepression. *A*’s setpoint is given by Eq 6. *A*_*env*_=0.2 m with a *K*_*i*_ of 7.735 × 10^−2^. The *k*_1_, *k*_2_ values are indicated in the figure. All other rate constants are as in Fig 4. Initial concentrations: *A*=0.696 m, *E*=0.018 au, *A*_*MRC*_=2.0×10^−3^ au, 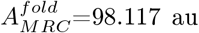. See Fig5.py for a corresponding Python calculation.

The term ‘aggressiveness’ relates to how fast a feedback responds towards a perturbation and how fast a controlled variable is moved to its setpoint. As an illustration for altered aggressiveness, *k*_3_ and *k*_5_ are changed by a common factor. This will influence the response kinetics of the NKA loop, but keep the setpoint of *A* unaffected. The violet curve in Fig 6 shows the response of *A* and *E* during two phases when at t=20 time units *k*_1_ is changed from 0.16 (phase 1) to 0.32 (phase 2). During both phases *k*_3_ and *k*_5_ are kept constant at respectively 1 × 10^−2^ and *k*_5_=1 × 10^−3^. This results in a smooth unimodal response of *A*, which takes around 180 time units to reach again *A*_*set*_=0.696 m.

**Figure 6:**
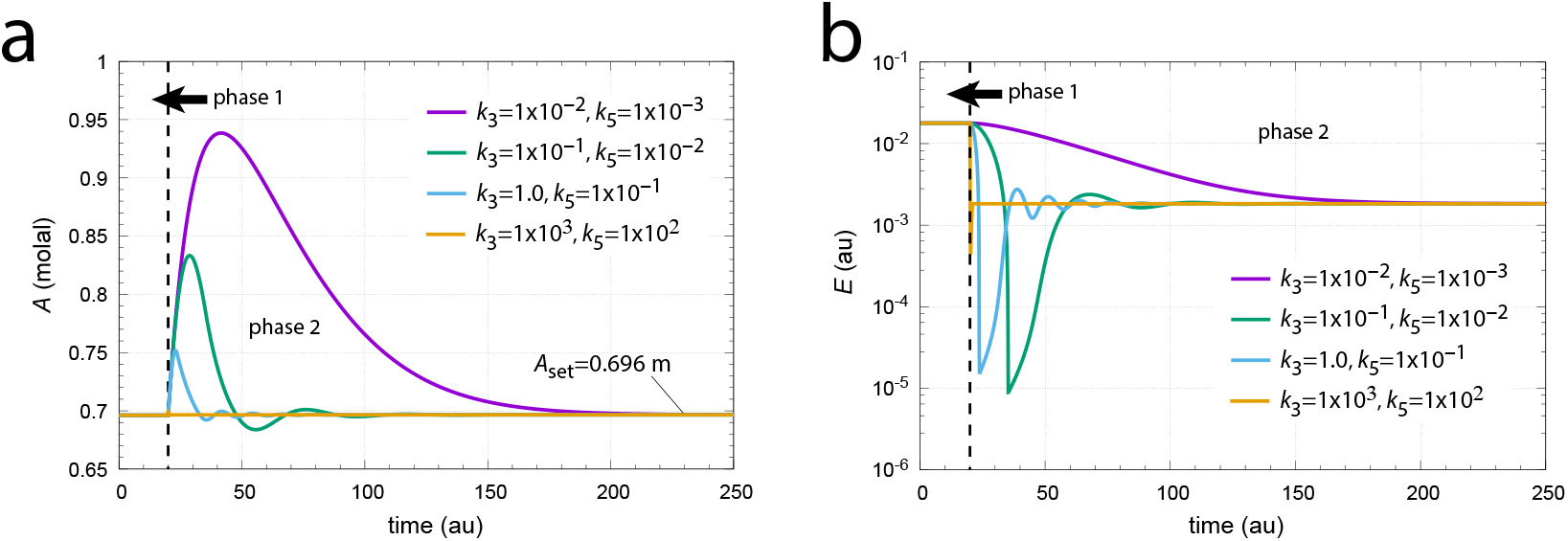
Illustration of NKA aggressiveness by changing *k*_3_ and *k*_5_ with a common factor. Panel a: Response of *A* to the step perturbation *k*_1_: 0.16 → 0.32 applied at *t* = 20 time units using four different *k*_3_, *k*_5_ combinations (see insets). As *k*_3_ and *k*_5_ increase *A*’s response becomes more aggressive. Panel b: The corresponding responses of *E*. See Fig6.py. for a corresponding Python script.

The green and blue lines show the effect when *k*_3_ and *k*_5_ are both increased by respectively one and two orders of magnitude. *A*’s adaptation becomes significantly faster, but is also accompanied by a damped oscillatory behavior. Finally, the orange line in Fig 6a shows the response when *k*_3_ and *k*_5_ are increased by five orders of magnitude each. The controller has become highly aggressive with a practically instantaneous adaptation to *A*_*set*_.

The ‘accuracy’ of a controller reflects its capability to adapt to the same setpoint independent of an applied perturbation. Fig 7 illustrates how 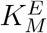 affects the NKA feedback’s accuracy. We have the same perturbations as in Fig 5, but 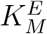 is now increased to 1×10^−3^. Controlled variable *A* no longer adapts perfectly to its previous setpoint (violet line), but the steady state values of *A* have become significantly dependent on the magnitude of the perturbation.

**Figure 7:**
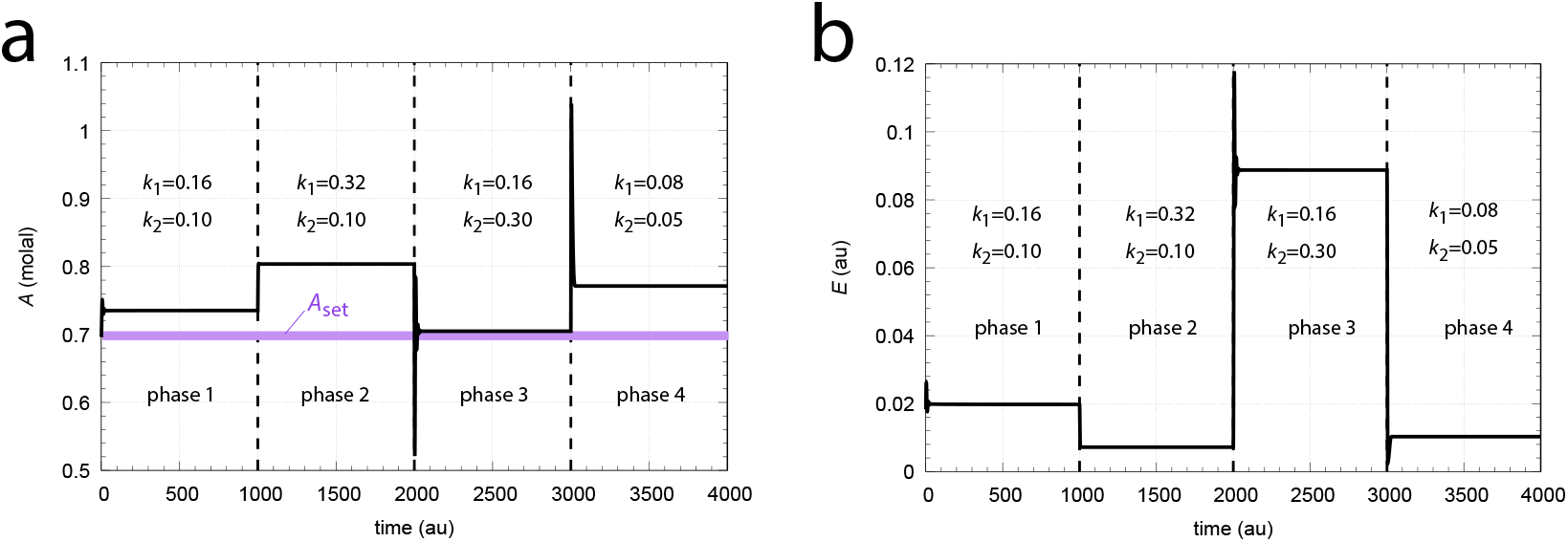
The accuracy of the NKA controller declines as 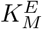 is increased. The system shown is the same as in Fig 5, but 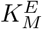 is increased to 1.0 × 10^−3^. *A*_*ss*_ now shows significant deviations above *A*_*set*_, which vary with the magnitude of the perturbation. See also Fig7.py.

The reason for this can be seen when considering a unit inhibition (*f*_*i*_(*A*)=1) in Eq 2. In this case the steady state of *E* is calculated as

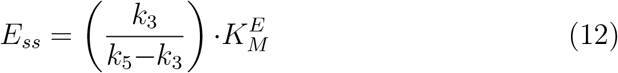

which is proportional to 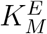. Thus, when 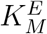 is increased also *E*_*ss*_ increases.This leads to larger compensatory sodium inflows into the blood compared with the situation when integral control is operative and is much lower. Due to the increased 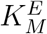 compensatory inflow at higher 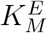, *A*_*ss*_ values will therefore stay above *A*_*set*_.

### 3.3. Other osmoregulatory schemes

Eqs 9, 10, and their resulting intersection point 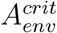 (Eq 11) determine the responses of the model, which include hyperosmotic as well as hyposmotic behaviors. This is indicated in Fig 8. Panels a and b show respective hyperosmotic and hyposmotic responses in *A* together with the corresponding repressions of *E* as *A*_*env*_ increases. The calculating procedure is the same as described for Fig 4. The changes in *K*_*i*_ affect *A*_*set*_ (Eq 6) prior to NKA break-down, while *k*_1_, *k*_2_, *a*, and *r* affect the 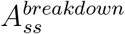 slope when *A*_*env*_ exceeds 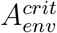.

**Figure 8:**
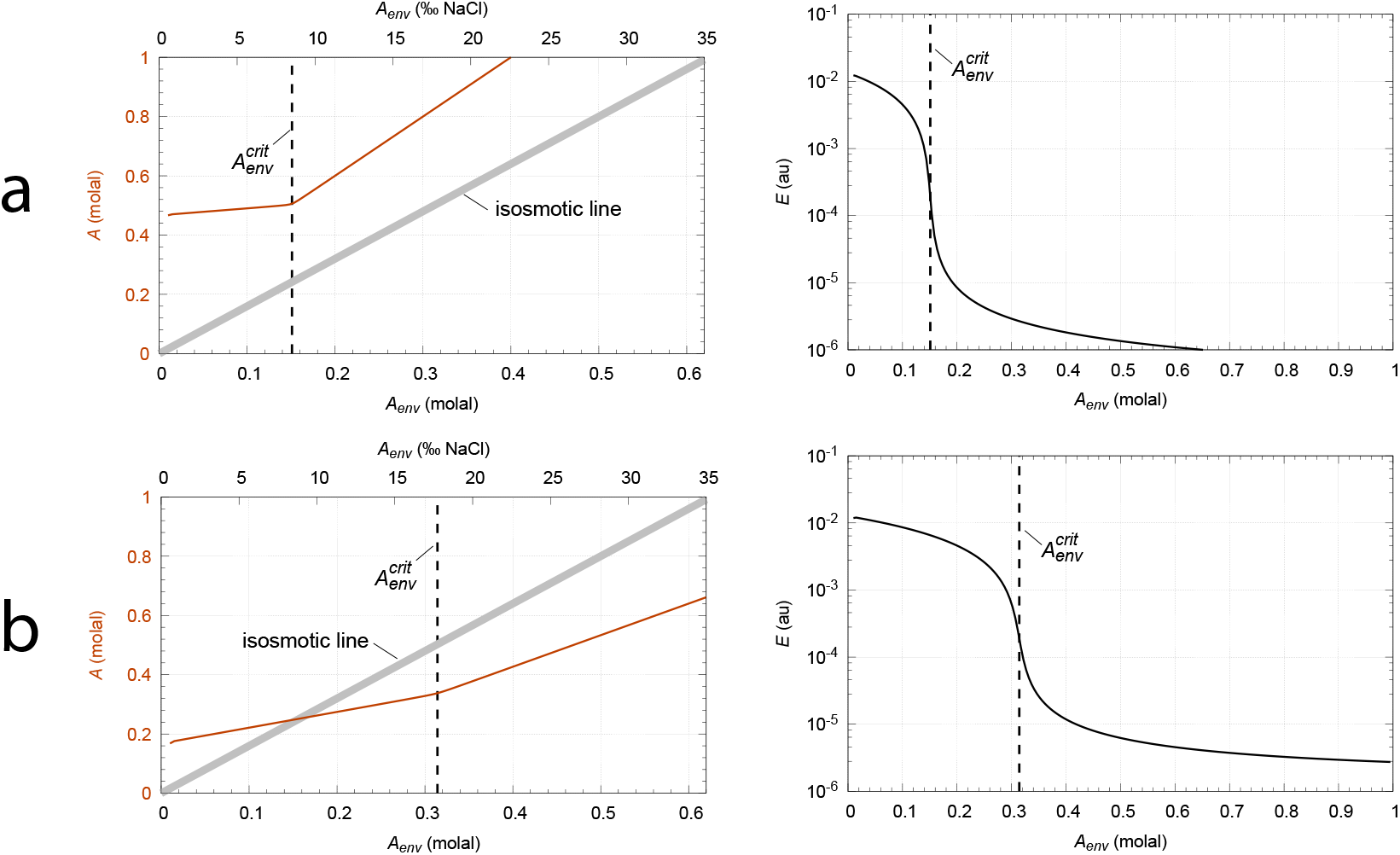
Hyperosmotic and hyposmotic responses of the model. Panel a: A hyperosmotic scheme where *A* does not cross the isosmotic line. Panel b: A hyposmotic response. *A* crosses the isosmotic line. Rate parameters panel a: 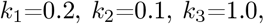 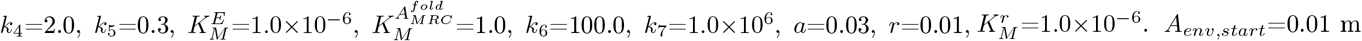 and *K*_*i,start*_=0.2 m, *A*_*env,end*_=1.0 m and *K*_*i,end*_=0.3 m. Initial concentrations, panel a: 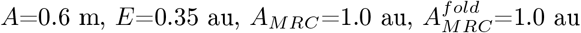 Rate parameters panel b: 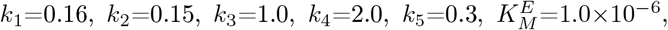 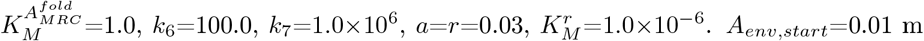 and *K_i,start_* = 0.072 m, *A_env,end_* = 1.0 and *K_i,end_* =0.3. Same initial concentrations as for panel a. For the Python scripts, see Fig8a.py and Fig8b.py.

## 4. Discussion

The term ‘enantiostasis’ was introduced by Mangum and Towle to distinguish the salinity adaptations of euryhaline animals from homeostatic behaviors (Mangum and Towle, 1977; Hochachka and Somero, 1984). The argument by Mangum and Towle assumed (as also done later by Mrosovsky (1990); Stevenson (2024) when introducing the concept of ‘rheostasis’) that homeostasis is related to a constant setpoint in relation with a single negative feedback loop. Although Cannon’s definition of homeostasis (Cannon, 1929) stressed the importance that steady states should be kept within narrow limits, Cannon didn’t claim that these steady states should be constant, but are the result of coordinated actions. In fact, Cannon seemed to be well aware about how crustaceans regulate the composition of their blood in dependency to environmental salinities. In his 1929 defining homeostasis paper he cited Fredericq (1885) on this matter. See further the comment by Langley (1973) how Fredericq’s work on regulation influenced Cannon’s view on homeostasis.

The model presented here is based on a single negative feedback loop by inhibiting NKA in parallel with the inflow of sodium ions into the blood via septate junctions. Excretion of sodium via urine and the sodium absorption *r* to various tissues enable a steady state in *A*. To keep the model as simple as possible, additional possible sodium outflows from the blood into the seawater via septate junctions (or otherwise) (Sáez et al., 2009; McNamara and Faria, 2012) have not been included as they serve the same function as sodium excretion.

MRCs undergo significant structural changes when environmental salinities vary (Mantel and Farmer, 1983; Luquet et al., 2002). These changes within MRCs may also lead to conformational changes of NKA, and possibly affect the enzyme’s rate parameters. For the sake of simplicity, it has been assumed that only *K*_*i*_ is affected by changed *A*_*env*_, while allosteric effects on the other rate parameters in Eq 6 are not considered.

As indicated in Fig 2b NKA is activated by internal sodium, while external sodium inhibits the enzyme. The reader may wonder what happened with the activation of NKA in the model Fig 3. Using shark NKA, Cornelius and Skou (1988) found that the Na^+^ half-maximal activation *K*_0.5_ was found to be about 6 mM, independent of the extracellular concentration of Na^+^. Thus, in the model I have simply absorbed the (assumed constant) activation term into *k*_3_.

Concerning the inhibition constant *K*_*i*_ of NKA by blood sodium, Sachs (1977) found, dependent on the formulations of the model, values between 4.4 mM and 12.0 mM. In our calculations *K*_*i*_ varied between 0.072 m and 0.099 m.

Interestingly, NKA can act both as an inflow or outflow controller, dependent on the enzyme’s transport orientation. Fig 9 gives a schematic overview. In panel a NKA acts as an outflow controller by moving the controlled species *A* out of the cell, while *A*_*env*_ is considered to be constant. The term *r*_*a*_ denotes the rate forming activated NKA^*^, which includes implicitly the inhibition by constant *A*_*env*_ and explicitly the activation by the controlled species *A*, i.e.

**Figure 9:**
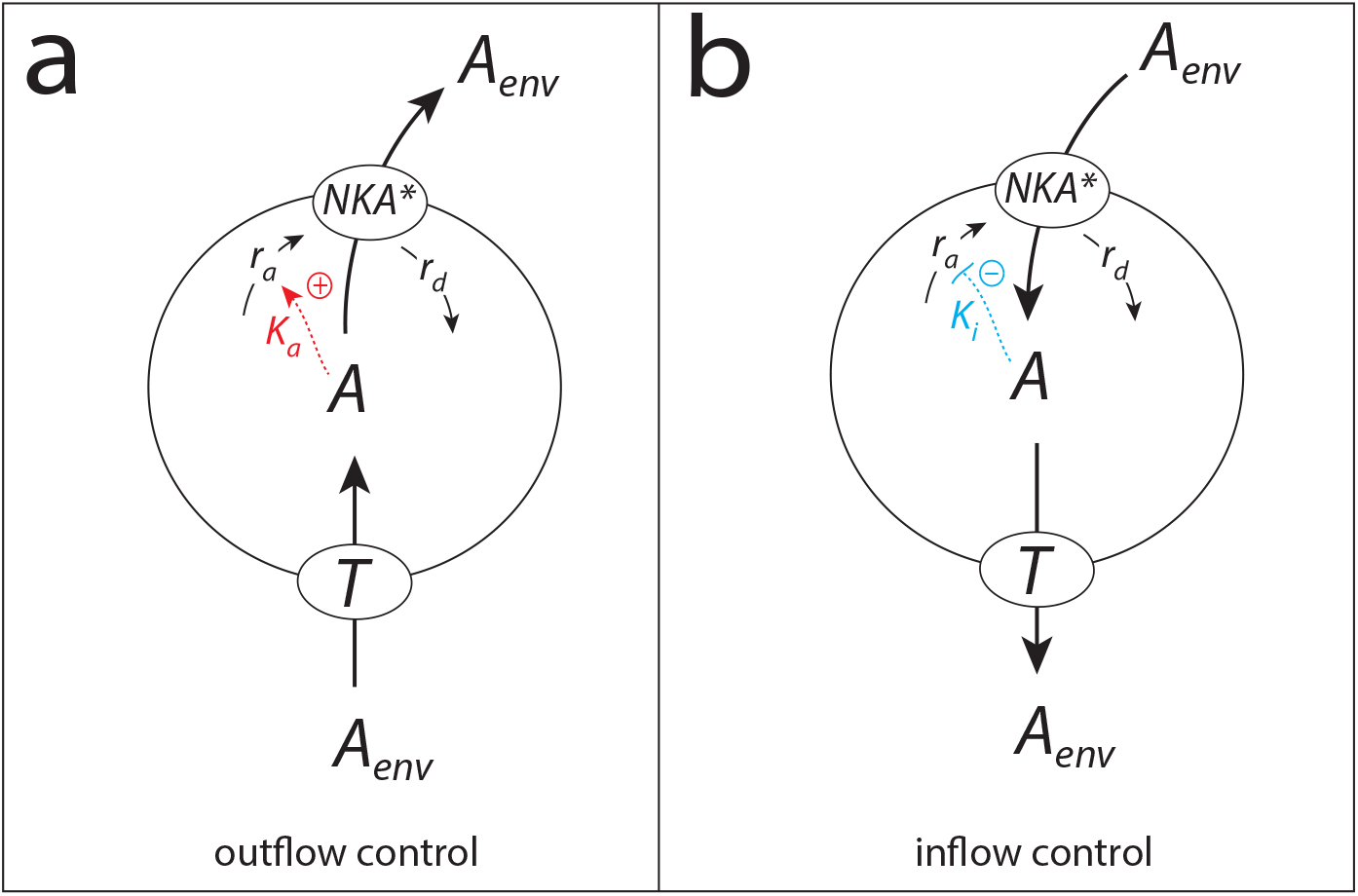
NKA can act as an outflow and inflow controller relative to the controlled variable *A*. Panel a: NKA acts as an outflow controller with an idealized integral control setpoint described by Eq 13. In panel b NKA acts as an inflow controller with setpoint described by Eq 14. NKA^*^ represents the activated pump.

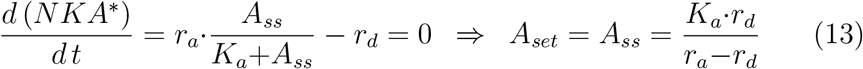

Fig 9b shows the inflow control by NKA as in Fig 3. The term *r*_*a*_ now implicitly contains NKA’s activation by *A*_*env*_ and explicitly the inhibition by the controlled variable *A*:

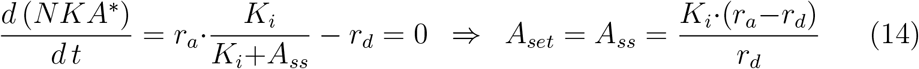

Thus, NKA may act in two ways dependent on the enzyme’s flux orientation. When acting as an outflow controller with respect to the controlled variable *A* (Fig 9a), we have a situation as observed for red cells where NKA moves sodium from the cell into the blood (Skou, 1998). Alternatively, when NKA acts as an inflow controller, as considered in this work, *A* is added to maintain homeostasis (Fig 9b).

A typical problem when dealing with simulations of biochemical processes is that experimentally determined rate parameters may not be available and that parameter values (to describe a certain outcome) depend on how a model is formulated. In our case many of the parameters used in the model are not known and possibly will depend on the type of organism or tissue. Furthermore, the model also represents an idealization when considering zero-order transitions from an activated NKA to the other NKA states in the Albers-Post cycle (Fig 2a). In fact, integral control may not be ideal since 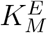 values may be larger; thus certain *A*_*env*_-based deviations from an ideal setpoint may occur as indicated in Fig 7 and in Drengstig et al. (2012). On the other hand, the data by Mangum and Towle (Fig 1a) appear to indicate a relative robust homeostatic mechanism where *A*_*set*_ increases with increasing environmental salinity. When comparing crabs treated in the laboratory (solid dots, Fig 1a) with crabs taken in the wild (open dots) a relative robust food homeostatic mechanism appears to be present, which indicates robustness in *A* regulation when model parameter *a* is varied.

Since Mangum and Towle’s introduction of enantiostasis (Mangum and Towle, 1977) several other concepts appeared as alternatives to Cannon’s homeostasis concept (Pereira Jr, 2021). They include rheostasis (Mrosovsky, 1990; Stevenson, 2024) and allostasis (Sterling and Eyer, 1988; McEwen and Wingfield, 2003) to mention two of them. They are often overlapping in their definition; for example they include setpoint changes, sometimes with the claim (Stevenson, 2024) that such changes cannot be explained by homeostatic mechanisms. As shown by the results from Figs 4 and 5 setpoint changes can indeed occur without compromising robust regulations. In this respect, behaviors addressed by enantiostasis or rheostasis, although important by their descriptions, appear to be emergent results of homeostatic regulation, either by single or multiple feedback loops.

## Supporting information

Python scripts for documentation

## Glossary

Controlled variable: The variable (here blood sodium *A*) which is under homeostatic control with one or several local setpoints
Controller breakdown: Controller breakdown occurs when the compensatory flux is insufficient, as for example when *j*_4_ (Eq 1) goes to zero
Controller variable or manipulated variable: The controller or manipulated variable generates a compensatory flux to oppose a perturbation applied to the controlled variable. In the NKA model this is the activated NKA variable *E*
Hyperosmosis: Osmolality of the blood is higher than the surrounding seawater
Hyposmosis: Osmolality of the blood is lower than the surrounding seawater
Integral control: In integral control the error between the controlled variable (or a function of it) and a local setpoint/reference is integrated and used as part of the negative feedback (see Eq 5). Integral control, when operative, ensures robust perfect adaptation to its setpoint when step perturbations are applied (see Fig 5)
Isosmotic line: The condition when osmolalities of blood and seawater are equal
NKA: Na-K ATPase, sodium pump

## Supporting Material

Python.zip: Python codes showing the results of Figs 4, 5, 6, 7, and 8. Besides Python (https://www.python.org), Numpy (https://numpy.org), Matplotlib (https://matplotlib.org), and Scipy (https://scipy.org) need to be installed. The scripts can be run via the Terminal (Linux, Unix, Mac OSX) or the Command Prompt (Windows) by using the command `‘python name.py’`, where `‘name’` is the name of the Python file.

