## Supplementary material for "Blood sodium regulation by Na,K-ATPase: Euryhaline animals’ salinity adaptations described by the pump’s negative feedback": Python scripts for documentation: Fig4.py

```
import numpy as np
import matplotlib.pyplot as plt
import time
from scipy.integrate import solve_ivp

# --- 1. Parameters ---
k1, k2, k3, k4, k5, k6, k7 = 0.16, 0.1, 1.0, 2.0, 0.1, 100.0, 1.0e6
ki_start, kme, kmamrc, a, r, kmr = 0.072, 1.e-6, 1.0, 0.022, 0.02, 1.e-6

Aenv_values = np.linspace(0.01, 1.0, 100) 
ki_values = np.linspace(0.072, 0.103, 100)
ki_slope = (0.103 - 0.072) / (1.0-0.01)

# critical Aenv calculation based on initial parameters
aenv_crit = (ki_start*((k3/k5)-1.0)-(a-r)/k2) / ((k1/k2)-ki_slope*((k3/k5)-1.0))

# --- 2. Define the ODE Model ---
def model_ivp(t, x, varAenv, current_ki):
    A, E, AMRC, AFOLD = x
    dAdt = k1*varAenv + k4*AFOLD*E/(kmamrc+AFOLD) - k2*A + a - r*A/(kmr+A)
    dEdt = k3*current_ki/(current_ki+A) - k5*E/(kme+E) 
    dAMRCdt = k6*varAenv - k7*AMRC
    dAFOLDdt = k7*AMRC - k4*AFOLD*E/(kmamrc+AFOLD) 
    return [dAdt, dEdt, dAMRCdt, dAFOLDdt]

# --- 3. Define the Jacobian (For Speed) ---
def jacobian_ivp(t, x, varAenv, current_ki):
    A, E, AMRC, AFOLD = x
    
    # Pre-calculate common denominators to save cycles
    den_E = (kme + E)**2
    den_AFOLD = (kmamrc + AFOLD)**2
    den_A_r = (kmr + A)**2
    den_A_ki = (current_ki + A)**2

    # Initialize a 4x4 zero matrix
    jac = np.zeros((4, 4))
    
    # Row 1: partials of dAdt
    jac[0, 0] = -k2 - (r * kmr) / den_A_r
    jac[0, 1] = (k4 * AFOLD) / (kmamrc + AFOLD)
    jac[0, 3] = (k4 * E * kmamrc) / den_AFOLD
    
    # Row 2: partials of dEdt
    jac[1, 0] = -(k3 * current_ki) / den_A_ki
    jac[1, 1] = -(k5 * kme) / den_E
    
    # Row 3: partials of dAMRCdt
    jac[2, 2] = -k7
    
    # Row 4: partials of dAFOLDdt
    jac[3, 1] = -(k4 * AFOLD) / (kmamrc + AFOLD)
    jac[3, 2] = k7
    jac[3, 3] = -(k4 * E * kmamrc) / den_AFOLD
    
    return jac

# --- 4. Define the steady state A event ---
def steady_state_event(t, x, varAenv, current_ki):
    derivs = model_ivp(t, x, varAenv, current_ki)
    return abs(derivs[0]) - 1e-9  # Focus only on A's steady state

steady_state_event.terminal = True
steady_state_event.direction = -1

# --- 5. Solve Loop with Timing ---
final_A, final_E = [], []
x0 = [0.6, 0.35, 1.0, 10.0] 

print("Starting solver...")
start_time = time.time()

for val_Aenv, val_ki in zip(Aenv_values, ki_values):
    sol = solve_ivp(
        model_ivp, (0, 2000), x0, 
        args=(val_Aenv, val_ki),
        method='LSODA',        # high-performance Fortran-based solver
        jac=jacobian_ivp,      # crucial for performance
        events=steady_state_event, 
        atol=1e-12, rtol=1e-9
    )
    
    # Extract steady state values
    ss_vals = sol.y[:, -1]
    final_A.append(ss_vals[0])
    final_E.append(ss_vals[1])
    
    # Update initial condition for the next step in Aenv
    x0 = ss_vals

end_time = time.time()
execution_time = end_time - start_time
print(f"Calculations finished in: {execution_time:.4f} seconds")

# --- 6. Plotting ---
aset_values = ki_values * ((k3 / k5) - 1.0)
isosmotic_line = 1.6 * Aenv_values
A_breakdown = (k1 * Aenv_values +a -r)/k2

fig, axs = plt.subplots(1, 2, figsize=(14, 6))

exp_Aenv, exp_A = np.loadtxt('data_MT.txt', unpack=True)
exp_A_corrected = exp_A / 1000.0
exp_Aenv_converted = exp_Aenv * (0.62 / 35)

# Panel 1: A vs Aenv
axs[0].plot(exp_Aenv_converted, exp_A_corrected, 'rs', 
            markerfacecolor='none', markeredgewidth=1.5, markersize=6, 
            label="Experimental ($A_{exp}$)", zorder=10)
axs[0].plot(Aenv_values, A_breakdown, color='orchid', lw=5, alpha=0.3, label="breakdown line")
axs[0].plot(Aenv_values, isosmotic_line, color='grey', lw=5, alpha=0.3, label="isosmotic line")
axs[0].plot(Aenv_values, aset_values, color='orange', lw=5, alpha=0.7, label="theoretical $A_{set}$")
axs[0].plot(Aenv_values, final_A, 'o', markersize=2, color='firebrick', label="numerical $A_{ss}$")
axs[0].axvline(aenv_crit, color='black', linestyle='--', alpha=0.8, label=r'$A_{env}^{crit}$')

axs[0].set_ylim(0.0, 1.1)
axs[0].set_xlim(0, 0.63)
axs[0].set_ylabel("$A_{ss}$ (molal)")
axs[0].set_xlabel("$A_{env}$ (molal)")
axs[0].legend(loc='upper left', fontsize='small')

# Secondary X-axis for salinity
secax = axs[0].secondary_xaxis('top', functions=(lambda x: x*(35/0.62), lambda x: x*(0.62/35)))
secax.set_xlabel('Salinity (‰ NaCl)')

# Panel 2: E vs Aenv
axs[1].plot(Aenv_values, final_E, color='black', lw=1.5)
axs[1].axvline(aenv_crit, color='black', linestyle='--', alpha=0.8)
axs[1].set_yscale('log')
axs[1].set_ylabel("$E$ (au)")
axs[1].set_xlabel("$A_{env}$ (molal)")

# Gridlines for both plots
for ax in axs:
    ax.minorticks_on() # Enables the small tick marks
    ax.grid(True, which='major', color='grey', linestyle='-', alpha=0.3)
    ax.grid(True, which='minor', color='grey', linestyle=':', alpha=0.1)

plt.tight_layout()
plt.savefig("Fig4_python.pdf", format='pdf', bbox_inches='tight')
print("Plot saved as Fig4_python.pdf")
plt.show()
```
