## Supplementary material for "Blood sodium regulation by Na,K-ATPase: Euryhaline animals’ salinity adaptations described by the pump’s negative feedback": Python scripts for documentation: Fig5.py

```
import numpy as np
import matplotlib.pyplot as plt
from scipy.integrate import solve_ivp

# --- 1. Parameters ---
Aenv, k3, k4, k5, k6, k7 = 0.2, 1.0, 2.0, 0.1, 10.0, 1.0e6
ki, kme, kmamrc, a, r, kmr = 0.0773487, 1.e-6, 1.0, 0.022, 0.02, 1.e-6

# Phase-specific k1 and k2
ks = [
    (0.16, 0.1), # Phase 1
    (0.32, 0.1), # Phase 2
    (0.16, 0.3), # Phase 3
    (0.08, 0.05)  # Phase 4
]

def model_ivp(t, x, vark1, vark2):
    A, E, AMRC, AFOLD = x
    dAdt = vark1*Aenv + k4*AFOLD*E/(kmamrc+AFOLD) - vark2*A + a - r*A/(kmr+A)
    dEdt = k3*ki/(ki+A) - k5*E/(kme+E) 
    dAMRCdt = k6*Aenv - k7*AMRC
    dAFOLDdt = k7*AMRC - k4*AFOLD*E/(kmamrc+AFOLD) 
    return [dAdt, dEdt, dAMRCdt, dAFOLDdt]                  

def jacobian_ivp(t, x, vark1, vark2):
    A, E, AMRC, AFOLD = x
    den_E = (kme + E)**2
    den_AFOLD = (kmamrc + AFOLD)**2
    den_A_r = (kmr + A)**2
    den_A_ki = (ki + A)**2

    jac = np.zeros((4, 4))
    # Row 1: dAdt
    jac[0, 0] = -vark2 - (r * kmr) / den_A_r
    jac[0, 1] = (k4 * AFOLD) / (kmamrc + AFOLD)
    jac[0, 3] = (k4 * E * kmamrc) / den_AFOLD
    # Row 2: dEdt
    jac[1, 0] = -(k3 * ki) / den_A_ki
    jac[1, 1] = -(k5 * kme) / den_E
    # Row 3: dAMRCdt
    jac[2, 2] = -k7
    # Row 4: dAFOLDdt
    jac[3, 1] = -(k4 * AFOLD) / (kmamrc + AFOLD)
    jac[3, 2] = k7
    jac[3, 3] = -(k4 * E * kmamrc) / den_AFOLD
    return jac

# --- 2. Simulation Setup ---
x_init = [6.9618E-01, 1.7818E-02, 2.0000E-03, 9.8117E+01]
t_all = []
y_all = []

current_x = x_init
t_start = 0

# --- 3. Loop through Phases ---
for i, (k1_val, k2_val) in enumerate(ks):
    t_span = (t_start, t_start + 1000)
    t_eval = np.linspace(t_span[0], t_span[1], 1000)
    
    sol = solve_ivp(
        model_ivp, t_span, current_x,
        args=(k1_val, k2_val),
        method='LSODA',
        jac=jacobian_ivp,
        t_eval=t_eval,
        atol=1e-12, rtol=1e-9
    )
    
    # Store results
    t_all.append(sol.t)
    y_all.append(sol.y)
    
    # Update for next phase: last column of sol.y
    current_x = sol.y[:, -1]
    t_start += 1000

# Concatenate all phases
t_final = np.concatenate(t_all)
y_final = np.concatenate(y_all, axis=1)

A_res = y_final[0, :]
E_res = y_final[1, :]
aset = np.full_like(t_final, 0.696)

# --- 4. Plotting ---
fig, ax1 = plt.subplots(figsize=(10, 6))

# Plot A on the left axis
ax1.plot(t_final, aset, color='orchid', linewidth=5.0, label='$A_{set}$=0.696')
ax1.plot(t_final, A_res, color='black', linewidth=1.5, label='A (blood sodium)')
ax1.set_xlabel("time (au)")
ax1.set_ylabel("$A$ (au)", color='black')
ax1.set_ylim(0.4, 1.1)

# Create the secondary axis for E
ax2 = ax1.twinx()
ax2.plot(t_final, E_res, color='blue', linestyle='--', alpha=0.7, label='E (activated NKA)')
ax2.set_ylabel("$E$ (au)", color='blue')

# --- Formatting the Right Ordinate to be Blue ---
ax2.spines['right'].set_color('blue')      # Changes the vertical line color
ax2.tick_params(axis='y', colors='blue')    # Changes the tick and label colors
# ------------------------------------------------

# Vertical lines for phase transitions
for phase_end in [1000, 2000, 3000]:
    ax1.axvline(x=phase_end, color='gray', linestyle='--', linewidth=2)

plt.title("Illustrating robust homeostasis in $A$ by varying $k_1$ and $k_2$")
ax1.grid(True, which='both', linestyle='--', alpha=0.5)

# Optional: Combine legends from both axes
lines1, labels1 = ax1.get_legend_handles_labels()
lines2, labels2 = ax2.get_legend_handles_labels()
ax1.legend(lines1 + lines2, labels1 + labels2, loc='upper right')

fig.tight_layout()
plt.savefig("Fig5_python.pdf")
plt.show()
```
