## Supplementary material for "Blood sodium regulation by Na,K-ATPase: Euryhaline animals’ salinity adaptations described by the pump’s negative feedback": Python scripts for documentation: Fig6.py

```
import numpy as np
import matplotlib.pyplot as plt
from scipy.integrate import solve_ivp

# --- 1. Parameters ---
# Please change the k3 and k5 values in the vector below to check the other Fortran results.
Aenv, k3, k4, k5, k6, k7 = 0.2, 1.e-2, 2.0, 1.e-3, 10.0, 1.0e6
ki, kme, kmamrc, a, r, kmr = 0.0773487, 1.e-6, 1.0, 0.022, 0.02, 1.e-6

# Phase-specific (k1, k2, duration)
# Phase 1: 20 units, Phase 2: 230 units
phases = [
    (0.16, 0.1, 20),  # (k1, k2, duration)
    (0.32, 0.1, 230), 
]

def model_ivp(t, x, vark1, vark2):
    A, E, AMRC, AFOLD = x
    dAdt = vark1*Aenv + k4*AFOLD*E/(kmamrc+AFOLD) - vark2*A + a - r*A/(kmr+A)
    dEdt = k3*ki/(ki+A) - k5*E/(kme+E) 
    dAMRCdt = k6*Aenv - k7*AMRC
    dAFOLDdt = k7*AMRC - k4*AFOLD*E/(kmamrc+AFOLD) 
    return [dAdt, dEdt, dAMRCdt, dAFOLDdt]                   

def jacobian_ivp(t, x, vark1, vark2):
    A, E, AMRC, AFOLD = x
    den_E = (kme + E)**2
    den_AFOLD = (kmamrc + AFOLD)**2
    den_A_r = (kmr + A)**2
    den_A_ki = (ki + A)**2

    jac = np.zeros((4, 4))
    jac[0, 0] = -vark2 - (r * kmr) / den_A_r
    jac[0, 1] = (k4 * AFOLD) / (kmamrc + AFOLD)
    jac[0, 3] = (k4 * E * kmamrc) / den_AFOLD
    jac[1, 0] = -(k3 * ki) / den_A_ki
    jac[1, 1] = -(k5 * kme) / den_E
    jac[2, 2] = -k7
    jac[3, 1] = -(k4 * AFOLD) / (kmamrc + AFOLD)
    jac[3, 2] = k7
    jac[3, 3] = -(k4 * E * kmamrc) / den_AFOLD
    return jac

# --- 2. Simulation Setup ---
x_init = [6.9618E-01,1.7818E-02,2.0000E-03,2.0625E+03]
t_all = []
y_all = []

current_x = x_init
t_start = 0

# --- 3. Loop through Phases ---
for k1_val, k2_val, duration in phases:
    t_span = (t_start, t_start + duration)
    # 10 points per time unit for smoothness
    t_eval = np.linspace(t_span[0], t_span[1], int(duration * 10))
    
    sol = solve_ivp(
        model_ivp, t_span, current_x,
        args=(k1_val, k2_val),
        method='LSODA',
        jac=jacobian_ivp,
        t_eval=t_eval,
        atol=1e-12, rtol=1e-9
    )
    
    t_all.append(sol.t)
    y_all.append(sol.y)
    
    current_x = sol.y[:, -1]
    t_start += duration

# Concatenate all phases
t_final = np.concatenate(t_all)
y_final = np.concatenate(y_all, axis=1)

A_res = y_final[0, :]
E_res = y_final[1, :]
aset = np.full_like(t_final, 0.696)

# --- 4. Plotting (Separate Subplots with Log Scale for E) ---
fig, (ax1, ax2) = plt.subplots(2, 1, figsize=(10, 8), sharex=True)

# Plot A on the top subplot
ax1.plot(t_final, aset, color='orchid', linewidth=3.0, label='$A_{set}$=0.696', alpha=0.6)
ax1.plot(t_final, A_res, color='black', linewidth=1.5, label='A (blood sodium)')
ax1.set_ylabel("$A$ (au)")
ax1.set_ylim(0.65, 1.0)
ax1.legend(loc='upper right')
ax1.grid(True, linestyle='--', alpha=0.5)
ax1.set_title("Illustrating feedback aggressiveness. Change $k_3$ and $k_5$ on line 7 in file 'Fig6.py' to see the other results.")

# Plot E on the bottom subplot
ax2.plot(t_final, E_res, color='blue', linewidth=1.5, label='E (activated NKA)')
ax2.set_yscale('log') # Set scale to logarithmic
ax2.set_ylim(1e-6, 1e-1) # Set limits from 1e-6 to 1e-1
ax2.set_xlabel("time (au)")
ax2.set_ylabel("$E$ (au) - Log Scale")

# Optional: Set specific ticks for clarity
ax2.set_yticks([1e-6, 1e-5, 1e-4, 1e-3, 1e-2, 1e-1])

ax2.legend(loc='upper right')
ax2.grid(True, which='both', linestyle='--', alpha=0.3) # 'both' shows major and minor grid lines

# Vertical line for the single phase transition at t=20
for ax in [ax1, ax2]:
    ax.axvline(x=20, color='red', linestyle=':', linewidth=2)

fig.tight_layout()
plt.savefig("Fig6_python.pdf")
plt.show()
```
