## Supplementary figures and images for "Blood sodium regulation by Na,K-ATPase: Euryhaline animals’ salinity adaptations described by the pump’s negative feedback"

### Fig4_python.pdf

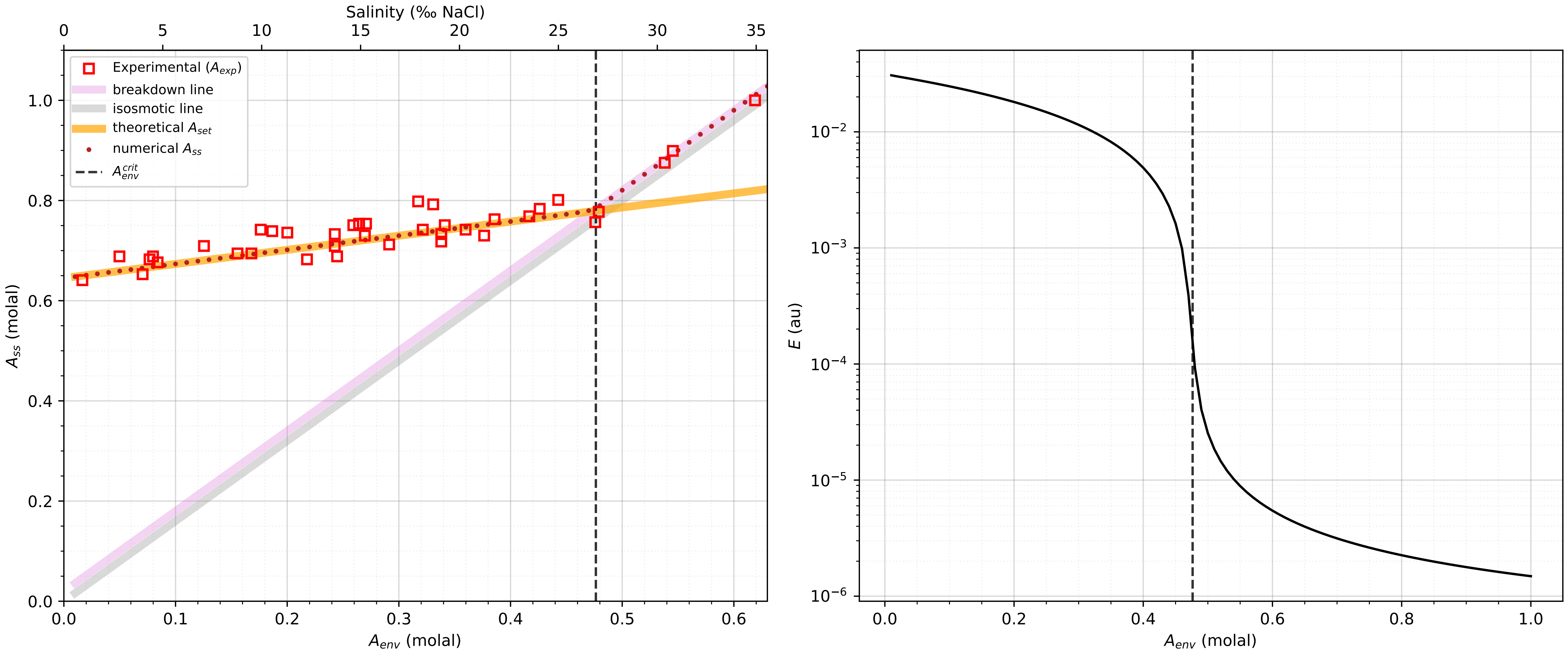

### Fig5_python.pdf

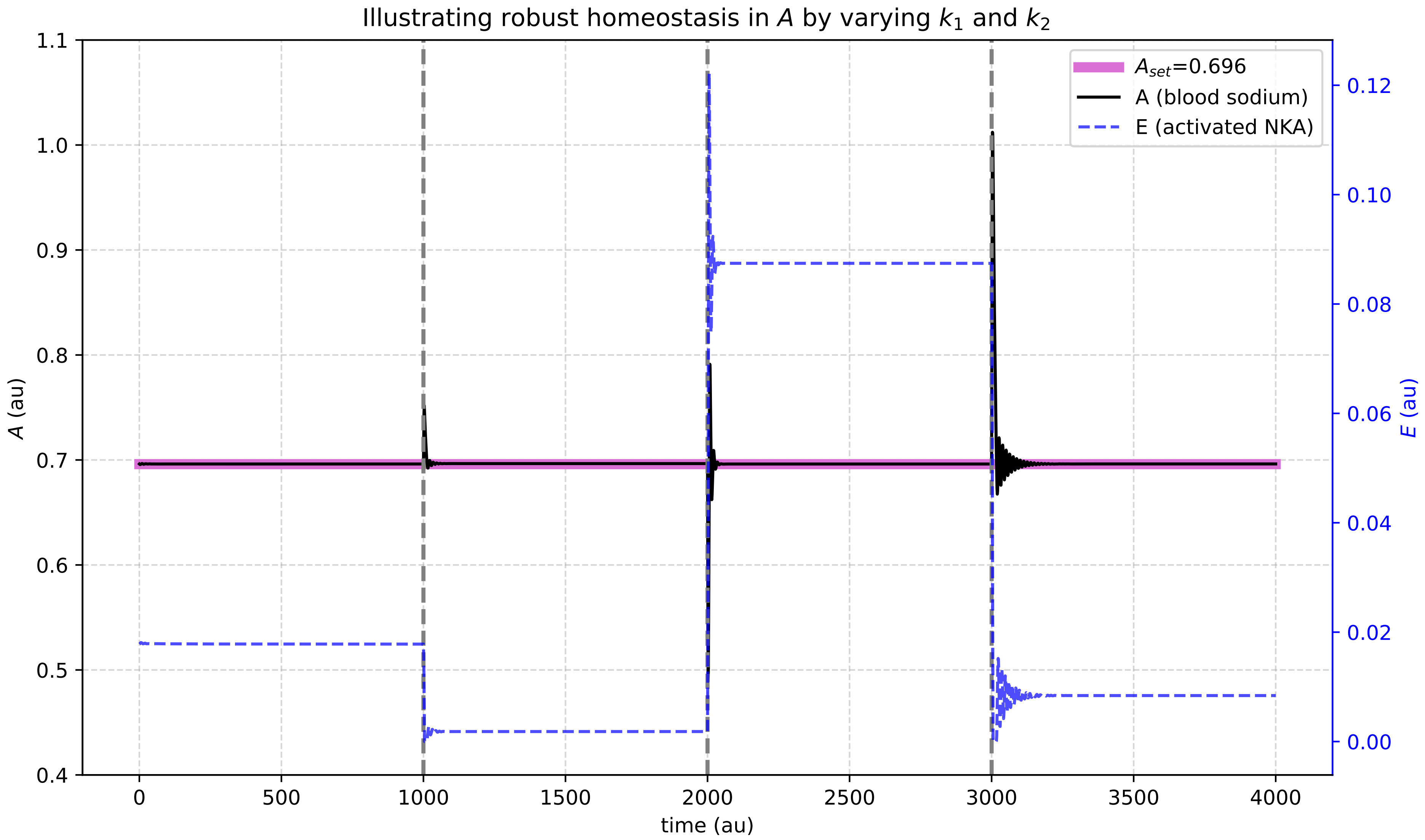

### Fig6_python.pdf

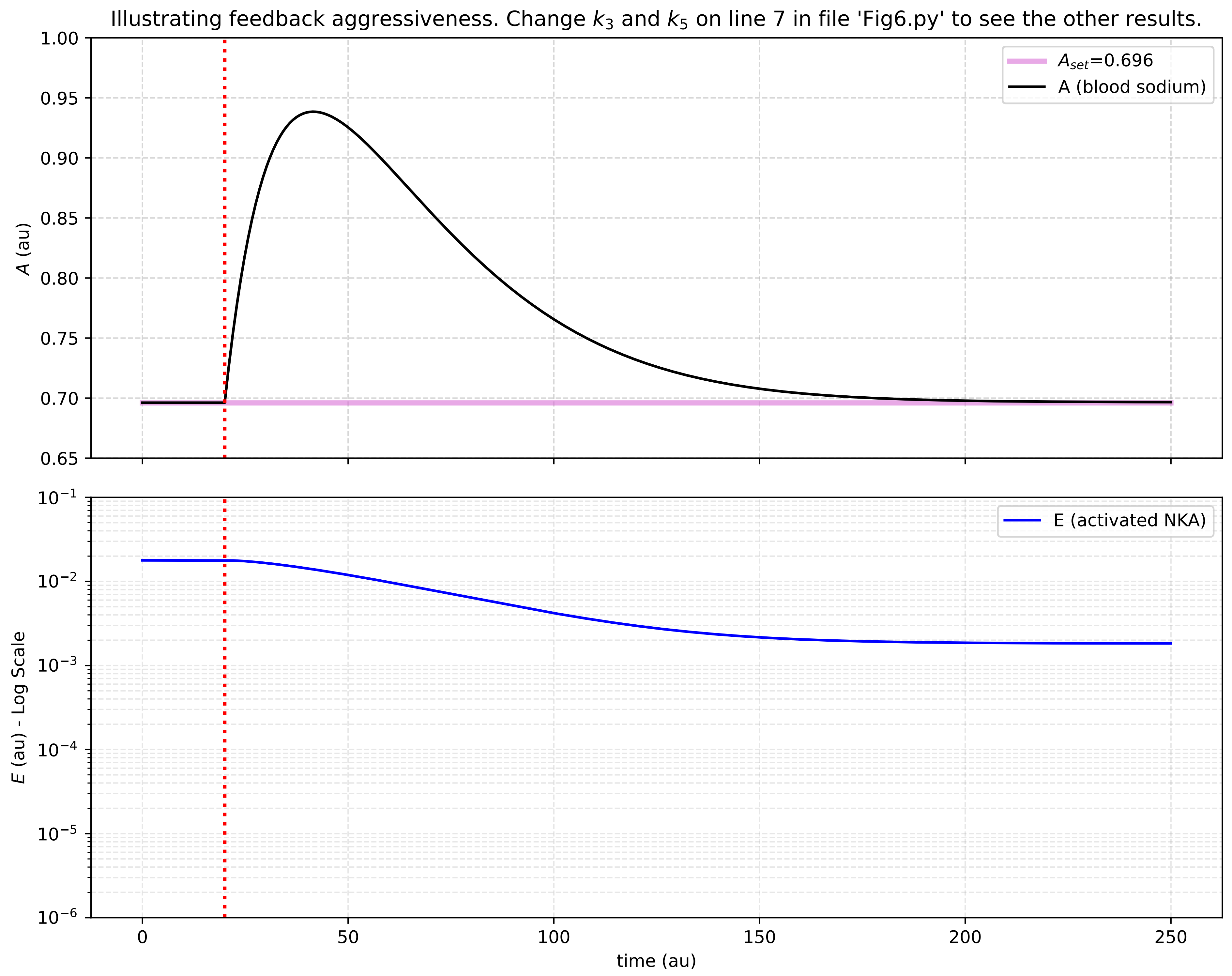

### Fig7_python.pdf

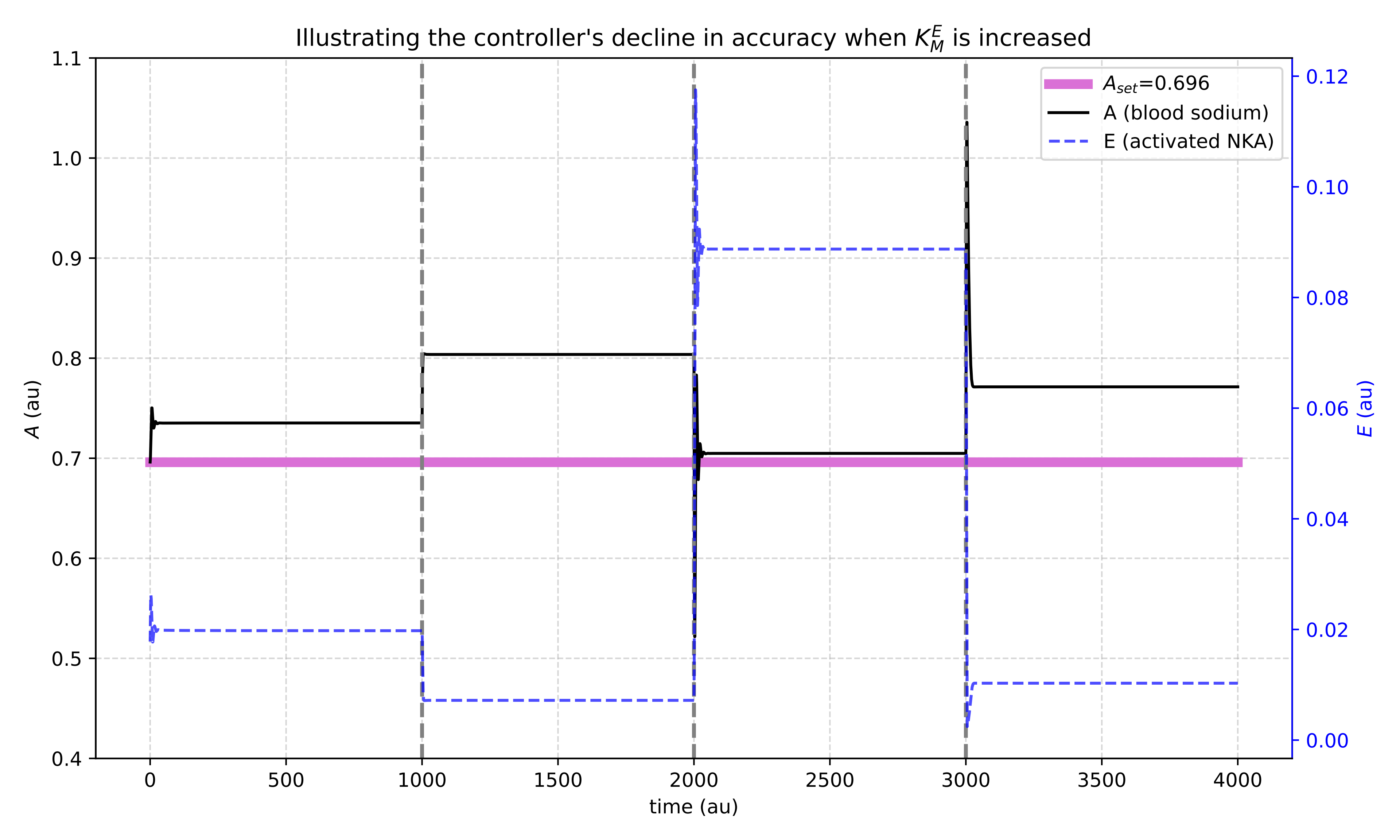

### Fig8a_python.pdf

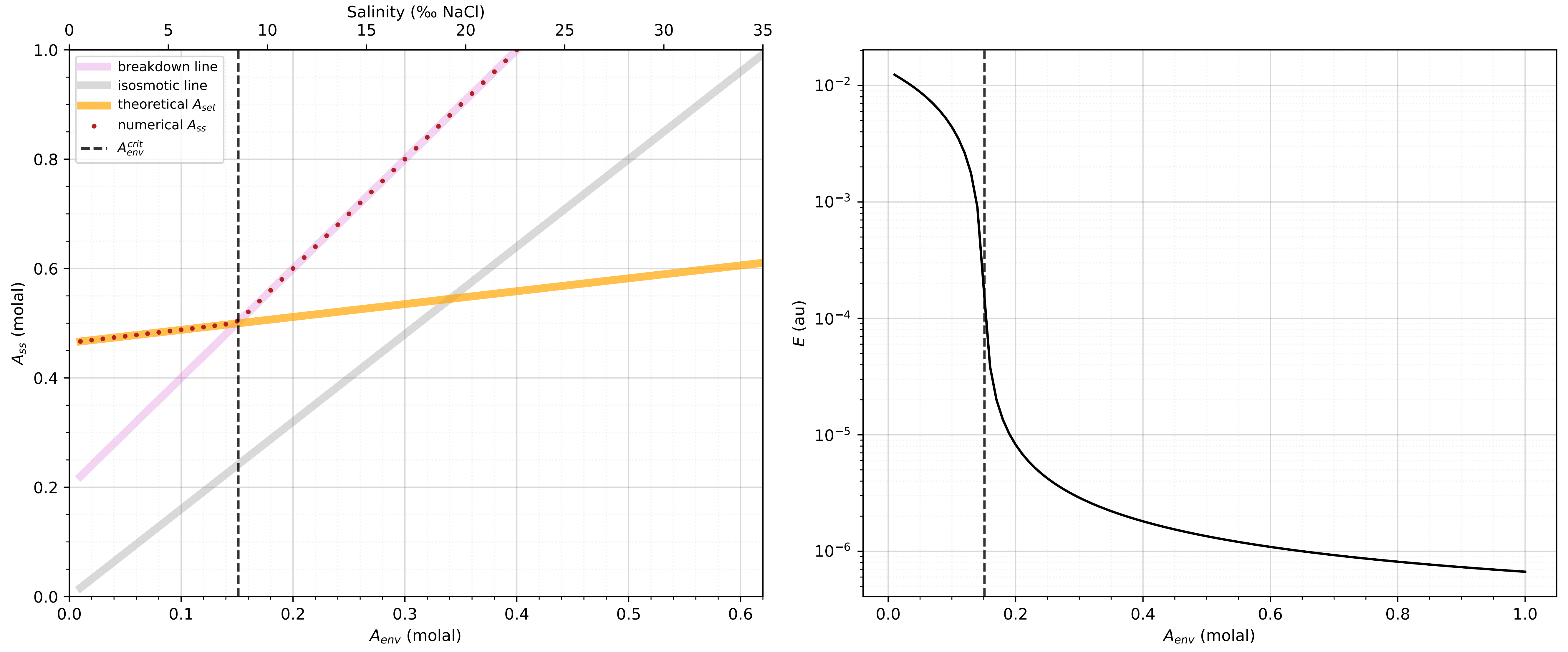

### Fig8b_python.pdf

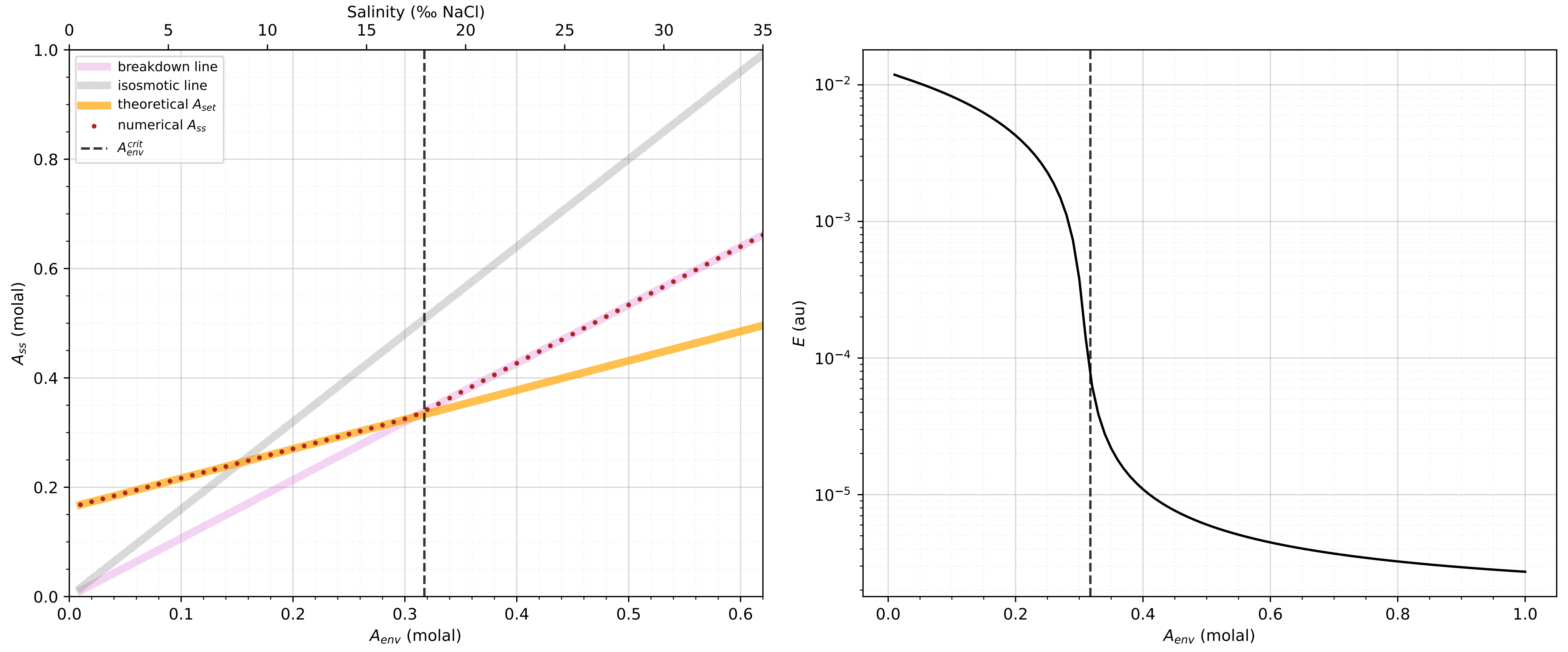
